# The effect of soil potassium and carbohydrates on xylem conductivity and embolism in an evergreen angiosperm tree and a gymnosperm tree before and after drought

**DOI:** 10.1101/2020.11.11.379156

**Authors:** Yael Wagner, Vlad Brumfeld, José M. Grünzweig, Tamir Klein

## Abstract

Xylem embolism is a major threat to tree function and survival under drought, in natural and agricultural settings alike, with its impact increasing in light of global climate change. Conversely, potassium (K^+^) has been shown to increase xylem conductivity (K_s_) in trees, and carbohydrates were reported to impact leaf gas exchange.

In this study we examined the effects of K^+^ and carbohydrates on K_s_ in two divergent evergreen tree species that are regularly exposed to drought: pine (*Pinus brutia*) and lemon (*Citrus × limon*). Five-year-old trees were pretreated with zero, moderate, and high K^+^, and with ambient or elevated CO_2_, to experimentally increase their xylem K^+^ or carbohydrates levels, respectively. Trees were then monitored for K_s_ and embolism (using a microCT), along with leaf gas exchange and water potential, before and after a 1.5-2.5 month drought period.

Potassium fertigation had a positive effect on K_s_, in both species when irrigated, which was eliminated following drought. Drought decreased K_s_ about 10-fold in lemon, with little effect in pine. CO_2_-treated trees had the same K_s_ as control trees before and after drought.

Our results indicate a positive effect of K^+^ on tree hydraulics, which was more pronounced in lemon than in pine, supporting the hypothesis of interaction with the angiosperm pit membrane, and not with the gymnosperm bordered pit. Yet, the elimination of this benefit following drought, and the lack of benefit from elevated carbohydrates following a short-term CO_2_ treatment, question the relevance of these components to tree drought resistance mechanisms.

**Key massage:** Potassium fertigation increases hydraulic conductivity and reduces xylem embolism in the gymnosperm pine, and more so in the angiosperm lemon tree, benefits which were eliminated following drought.

## Introduction

Global climatic change is believed to increase the future frequency and intensity of drought events (Solomon et al. 2007; McDowell et al. 2018). Greater variance in temperature and water availability will require trees in drier areas to perform under increasingly harsher drought-stress, thus to adapt effective drought resistance and recovery mechanisms (Solomon et al. 2007; Allen et al. 2010; Choat et al. 2012; Anderegg et al. 2016). In response to desertification processes already taking place in many areas and to a decrease in weather predictability (Solomon et al. 2007), as well as the increase in extreme precipitation and changes in seasonal precipitation (Zeppel et al. 2014b), understanding mechanisms enabling the prevalence of trees through drought events is crucial in predicting future trends in tree survival and distribution.

Drought-induced tree mortality is largely caused by hydraulic failure (McDowell et al. 2008, 2018, Anderegg et al. 2012, 2015, 2016; Adams et al. 2017; Choat et al. 2018). This happens when trees lose a substantial fraction of their hydraulic conductivity due to cavitation, when air bubbles are formed inside the xylem conduits (Pratt et al. 2008, Jacobsen et al. 2018). The formation of the bubbles takes place under dry environmental conditions, when water availability decreases. As a result, xylem tension increases (i.e. the water potential becomes more negative). At a certain threshold tension, pit membrane failure allows air from non-functioning vessel into a water-filled vessel in a process called ‘air-seeding’ (Pickard 1981; Wheeler and Stroock 2009; Rockwell et al. 2014). The blockage created by the bubbles is called embolism.

The complete set of traits and strategies trees incorporate to avoid hydraulic failure when operating under low water-availability is unknown. However, several factors have been shown to play a role in trees’ response to drought. For example, changes in ion concentration in xylem sap are known to affect hydraulic conductivity of plants (Zimmermann 1978; Zwieniecki et al. 2001). Ions have been known to influence xylem conductivity for decades (Zimmermann 1978). Potassium (K^+^) in particular has a great potential to affect such processes, as its abundance in the xylem sap is high compared to other ions (Herdel et al. 2001). High concentrations of ions in sap fluid have been shown to increase hydraulic conductivity by up to 60% (Zwieniecki et al. 2001; Nardini et al. 2011). However, the basic mechanism of this effect is still debated. One hypothesized mechanism is based on the interaction between ions in the sap and pectins, which are a major component of the pit membrane. The state of the pectins ranges from gel to amorphous solid as a function of their hydration status (Ryden et al. 2000). High cation concentration in the sap fluid increases the hydration rate of the pectic matrix in the pit membrane and causes its shrinkage (Zwieniecki et al. 2001; Lee et al. 2012), consequently decreasing the resistance it poses to water flow. Another hypothesis is based on the electro-viscous effect (Kirby 2010), where the water flowing through the pits wash out ions from their surface and create an electrical gradient in the opposite direction to the water potential gradient. Highly deionized water would wash the ions more readily and create higher electrical gradient, which would, in turn, slow down the flow through the pit and increase its hydraulic resistance (Santiago et al. 2013). Whichever is the underlying mechanism for the so-called ‘ion-effect’, it seems as though pit-membrane structure would have a key impact on its magnitude. In consistent with that, an ion effect was found to increase the conductivity of the membrane and consequentially the hydraulic conductivity of the plant, mainly in angiosperms (van Ieperen et al. 2000, Zwieniecki et al. 2001, Zwieniecki et al. 2003, Lopes-Portillo et al. 2005, Trifilo et al. 2008, Aasamaa and Sõber 2010, Nardini et al. 2011). Additionally, drought stressed trees were found to buffer loss of hydraulic conductivity through increase in sap K^+^ levels compared to non-stressed ones (Trifilò et al. 2011). However, when measuring drought recovery, Oddo et al. (2014) found no difference between trees that were irrigated with KCl solution and trees that received K^+^ free water. Therefore, the complete picture of the role of K^+^ in plant hydraulic conductivity is not clear.

Another group of compounds that were hypothesized to play a role in the dynamics of tree hydraulic conductivity is carbohydrates, which are assumed to take part in several processes supporting tree survival under drought. Some of these processes can be studied by manipulating carbon availability at the whole-tree level, e.g. via CO_2_ enrichment. Atmospheric CO_2_ levels can affect the tree’s hydraulic state in several ways. First, higher concentration of CO_2_ could lead to higher influx into the mesophyll tissue, increasing CO_2_ concentration in chloroplasts and turning photosynthesis more efficient (Gaastra 1959). This would potentially allow earlier stomatal closure, which may, in turn, reduce the amount of water lost through transpiration (Paudel et al. 2018). Alternatively, in Eucalyptus, elevated CO_2_ was found to promote shallower rooting depth and increased leaf area, which may lead to increased drought stress (Duursma et al. 2011). Second, increased CO_2_ levels may lead to an increase in stored carbohydrates in the tree (Paudel et al. 2018; Li et al. 2018; Duan et al. 2018). Trees that have higher levels of stored carbohydrates may survive longer periods with closed stomata and low physiological functioning without risking carbon starvation (Klein et al. 2014).

Clarifying the role of carbohydrates and K^+^ in trees’ ability to survive drought events is one step towards completing our view of the full set of mechanisms allowing trees to maintain hydraulic functionality under low water availability. In the context of rising temperatures and decrease in precipitation, this knowledge is crucial for the prediction of future trends in tree mortality and distribution, as well as for the development of appropriate agricultural and forestry practices. In this study we examined two evergreen tree species that are regularly exposed to drought. Lemon (*Citrus × limon*) is a cultivated angiosperm species originated in south-east Asia (Zohary et al. 2012) and is being commercially grown all around the Mediterranean basin. It typically receives no irrigation for two months during the summer (July-August) as an agricultural practice to induce autumnal flowering and an off-season yield (Raveh 2008). Turkish pine (*Pinus brutia*) is a coniferous forest tree species typically found in East-Mediterranean habitats, where water availability is the main limiting factor to growth (Zohary 1966). We studied the effect of different levels of K^+^ and atmospheric CO_2_ on the ability of young (five years old) potted lemon and pine trees to withstand the hydraulic stress induced by a 1.5-2.5 month of drought (for lemon and pine, respectively). The trees were treated with one of three levels of K^+^ to test its effect on hydraulic conductivity, and one of two levels of air-CO_2_, to experimentally increase their sap soluble sugar levels, for a month prior to being dried. This is opposed to a post-treatment addition of potassium, which showed no effect on hydraulic recovery (Oddo et al. 2014). Tree hydraulic and physiological state were monitored before and after the drought period, in addition to their sap K^+^ and sugars levels. We hypothesized that (1) high K^+^ will increase K_s_ before, and potentially also after, drought; and (2) high sugar will decrease embolism level (and thereby increase K_s_) during recovery from drought.

## Materials and methods

### Experimental design and plant material

The experiment was conducted in the Gilat CO_2_ enrichment greenhouse (31°33”N; 34°66”E) and at the Weizmann Institute greenhouse (31°90”N; 34°80”E). The light, temperature and humidity in both locations were coupled to the local ambient conditions. We used five-years-old trees of Lemon (*Citrus × limon* cv. Eureka) and Turkish Pine (*Pinus brutia*) (20 from each species). Lemon and pine tree height was 210 ± 3 and 220 ± 3 cm, respectively; stem diameter at base was 20.98 ± 0.34 and 22.54 ± 0.88 mm, respectively. Trees were potted in 35 L pots filled with inert perlite media, were exposed to one of two levels of atmospheric CO_2_ (400 and 850 ppm) and received one of three levels of potassium fertigation (0, 40 and 80 ppm) during a month, a period termed here the ‘pre-drought treatment’ (Fig. 1). Trees that received 400 ppm CO_2_ and 40 ppm K^+^ acted as ‘control’ trees. In total, 20 trees of each species were used in a non-matrix design, since treatment combinations were not the focus of this study, but rather independent effects of potassium levels and CO_2_-induced sugar loading. For the complete fertilization formulas, see Table S1. The CO_2_ levels were imposed by continuously pumping CO_2_ into one room in the greenhouse to create concentration of 850 ppm while another room was kept at ambient conditions and monitored to avoid the formation of excess CO_2_ levels. Temperatures, humidity and radiation levels were monitored in both rooms and did not vary between them. During the pre-drought treatment, midday temperature in Gilat greenhouse ranged from 20°C to 30°C and relative humidity from 45% to ~90%. Next, trees were exposed to drought (i.e. zero irrigation) for 40 (lemon) or 74 (pine) days at the Weizmann greenhouse, where ambient midday temperature increased gradually with seasonal change from 10-20°C during all of the lemon drought period to 20-25°C at the end of pine drought period. Relative humidity varied, and was averaged around 50% throughout drought periods. Four campaign days were conducted, one for each species at the end of the pre-drought treatment (and prior to the drought onset) and one for each species at the end of the drought period, during which hydraulic conductivity and leaf gas exchange were measured for three trees from each treatment.

**Fig. 1.**
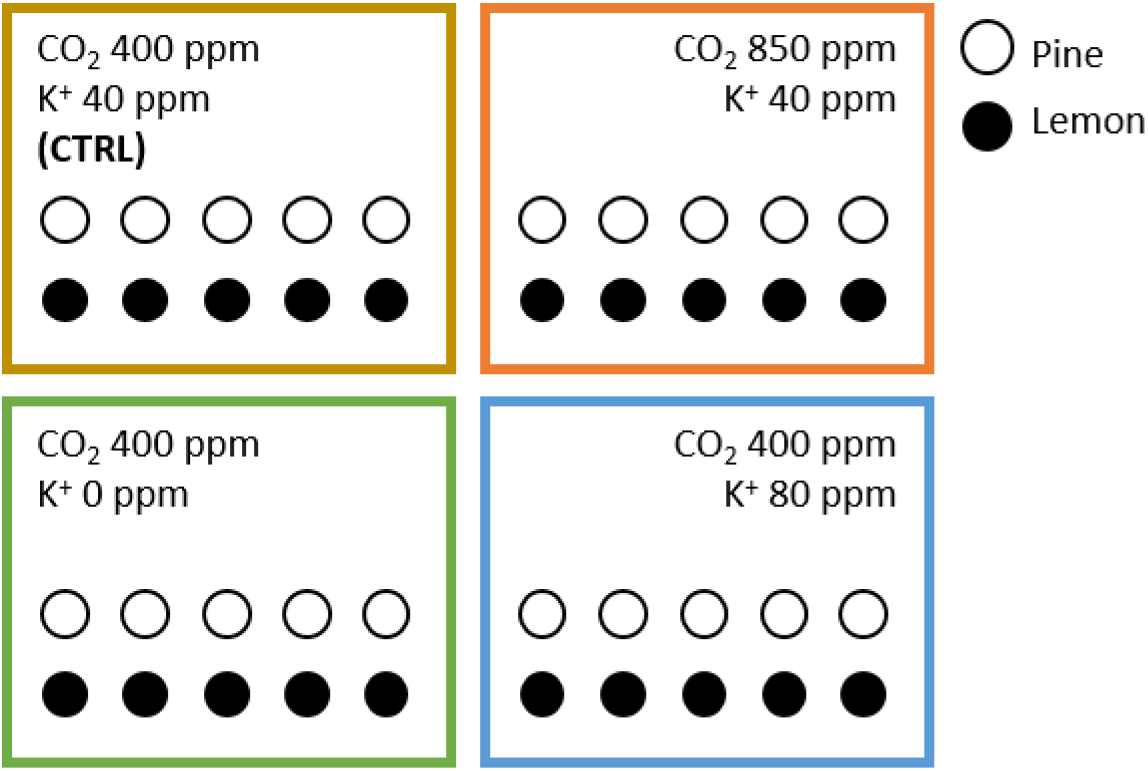
Experimental layout.

### Xylem hydraulic conductivity

Xylem specific hydraulic conductivity under native conditions (K_s_) was measured at drought onset and at the end of drought treatment according to a protocol adjusted from Sperry et al. (1988) and Wheeler et al. (2013). The measurements were carried out on three trees from each species and treatment. Branches (~40 cm long) were cut from the trees under degassed water and submerged in water immediately (with pine branches, ice water was used to stop resin from oozing and blocking the xylem conduits) in order to prevent the inducement of artificial embolism. The branches were kept under water for ~30 minutes to allow the relaxation of xylem tension and were then further cut under water into 10 cm segments. The final segments were placed under a column of degassed 19 mM KCl solution with hydrostatic pressure of 0.7 kPa. This solution is standard for hydraulic measurements across numerous studies, and was used here across all measurements, regardless of the potassium treatment. During sampling, all precautions were taken and comparisons were only conducted within species in order to avoid biased results. Vessel length distribution was measured in 0.9 m long branches sampled from four 20-years-old lemon trees grown on campus in 2016 (Cohen & Paudel, unpublished data). These measurements used the improved model developed by Cohen et al. (2003) based on air injection and conductivity measurements. The measured skewed distribution function peaked at ~6 cm, with only 5% of the vessels being 20 cm long, and 1% at 40 cm. Here we cut 40 cm branch segments under water, from which 10 cm segments were analyzed. Considering the vessel length distribution, the fraction of open vessels among the air-filled vessels in lemon xylem was minimal. Another evidence for this is the very low number of air-filled vessels in potassium-fertigated lemon before the drought (see results).

The hydraulic conductivity (K_h_) and sapwood specific hydraulic conductivity (K_s_) of the branch segments were finally calculated as follows:

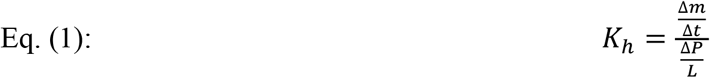

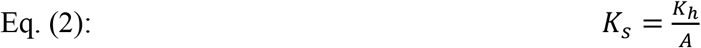

Where K_h_ is the hydraulic conductivity (kg m sec^−1^ MPa^−1^); K_s_ is the specific hydraulic conductivity (kg s^−1^ m^−1^ MPa^−1^); Δm is the mass of water conducted by the branch segment (kg); Δt is the time interval for the measured conductance (sec); ΔP is the change in hydrostatic pressure created because of water conductance and decrease of the water column (MPa); L is the length of the measured branch segment (m), and A is the cross section of the conducting area (m^2^).

### MicroCT imaging

MicroCT scans were performed on branch segments of one tree from each species and treatment using an X-radia microCT located in the Weizmann Institute (Zeis X-ray microscopy, Peasanton, USA). Segments were taken from one of the branches used to measure hydraulic conductivity (one segment per treatment). Each segment was wrapped in Parafilm and placed in the microCT. We took 700 projections over 180° rotation range, using optical magnification of 0.5x in order to increase the field of view. Voltage and current were set to 30 kV and 150 μA, respectively. The resulted voxel size ranged between 8.0-9.6 μm. Volume reconstruction was performed by a filtered back-projection algorithm (Zeiss X-ray microscopy, Peasanton, USA). The 3D model of the branch, shows dark regions representing the least dense tissue (i.e. air) and bright regions that represent the densest tissues (i.e. minerals, meristematic tissues). Xylem tissues like conduit walls, parenchyma, fibers etc. appear in the image having different shades of grey.

### Leaf water potential and gas exchange

Leaf water potential (Ψ_1_) was measured using a pressure chamber (PMS Instrument Company, New York, USA) every two weeks on one tree from each species and treatment and during campaign days on all trees used for hydraulic conductivity measurements. Leaves were sampled from the trees (branch tips in pine) and kept in sealed, foil-covered bags on ice for a maximum of ten minutes before they were brought to the lab and measured. Assimilation (A), transpiration (E) and stomatal conductance (g_s_) were measured using an infra-red gas analyzer (WALZ GFS-3000, Germany; LiCor6400XT, Nebraska, USA) during campaign days on all the trees used for hydraulic measurements. Leaves or ten pairs of needles were sampled from lemon and pine trees, respectively, and immediately put into the gas exchange system cuvette, where constant CO_2_ level of 400 ppm was maintained and light was adjusted to ambient. Measurements were performed three times along the day: morning (08:00-10:00), midday (12:00-14:00) and evening (16:00-18:00). Analyses were performed on diurnal means, as well as on discrete time-points.

### Statistical analysis

Results of leaf gas-exchange, leaf water potential and hydraulic conductivity from each species and campaign day were pooled, and subjected to two-way ANOVA using JMP^®^ software (Cary, NC, USA), where the factors were treatment (4 levels) and date (2-5 levels) The results of the ANOVA are reported in Table 1. The data were further analyzed for means comparison using Student’s *t* test. Our analyses are based on diurnal averages of morning, midday, and evening measurements.

**Table 1.**
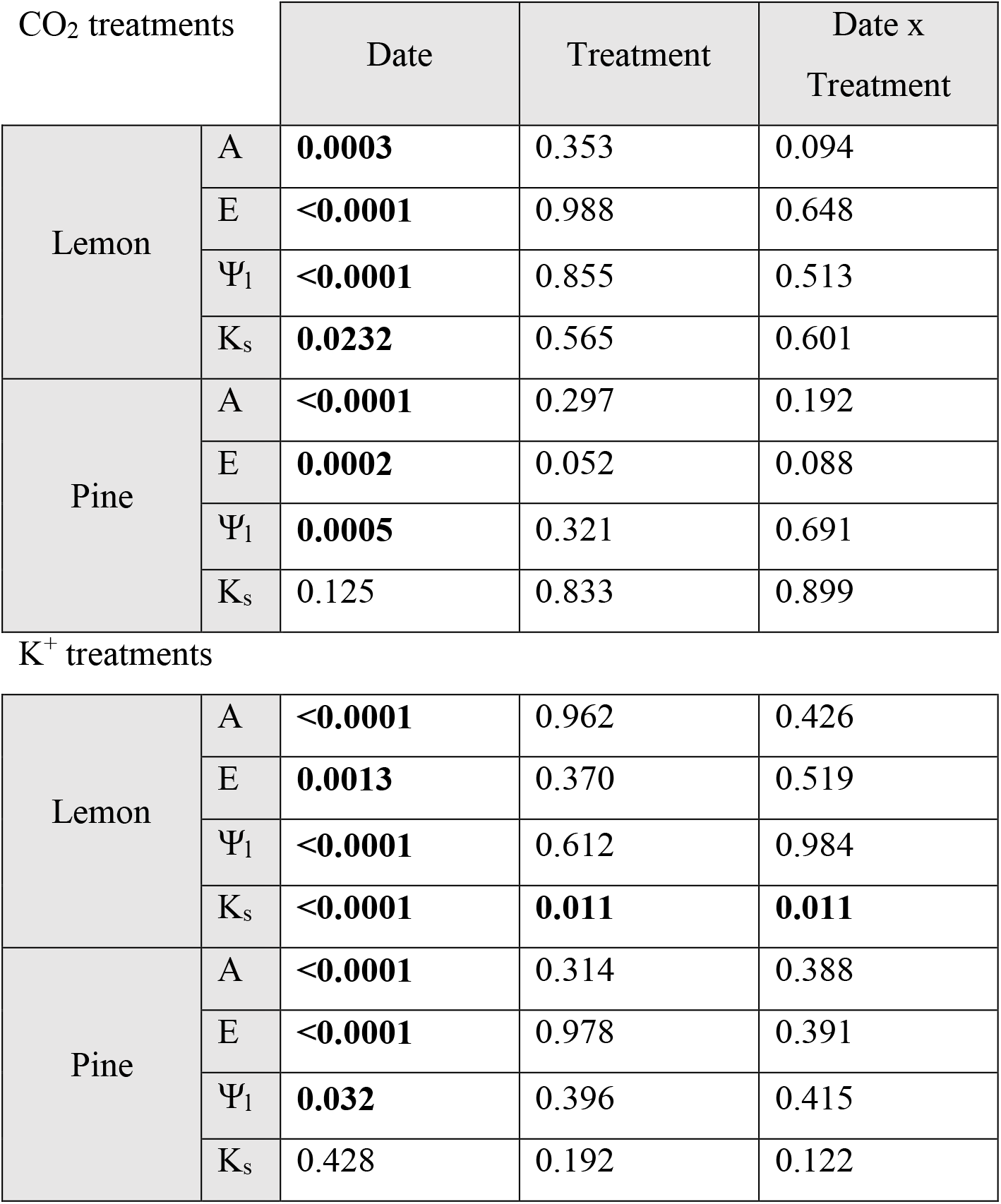
*P*-values obtained from two-way ANOVA for the effects of date (drought onset, drought end), treatment, and their interaction, on four plant responses: A, assimilation; E, transpiration; Ψ_1_, leaf water potential and K_s_, xylem hydraulic conductivity. Species were analyzed separately. Significant effects appear in **boldface**.

| CO<sub>2</sub> treatments |  | Date | Treatment | Date x Treatment |
| --- | --- | --- | --- | --- |
| Lemon | A | **0.0003** | 0.353 | 0.094 |
|  | E | **<0.0001** | 0.988 | 0.648 |
| | $\Psi_l$ | **<0.0001** | 0.855 | 0.513 |
| | $K_s$ | **0.0232** | 0.565 | 0.601 |
| Pine | A | **<0.0001** | 0.297 | 0.192 |
|  | E | **0.0002** | 0.052 | 0.088 |
| | $\Psi_l$ | **0.0005** | 0.321 | 0.691 |
| | $K_s$ | 0.125 | 0.833 | 0.899 |
| K<sup>+</sup> treatments |  |  |  |  |
| Lemon | A | **<0.0001** | 0.962 | 0.426 |
|  | E | **0.0013** | 0.370 | 0.519 |
| | $\Psi_l$ | **<0.0001** | 0.612 | 0.984 |
| | $K_s$ | **<0.0001** | **0.011** | **0.011** |
| Pine | A | **<0.0001** | 0.314 | 0.388 |
|  | E | **<0.0001** | 0.978 | 0.391 |
| | $\Psi_l$ | **0.032** | 0.396 | 0.415 |
| | $K_s$ | 0.428 | 0.192 | 0.122 |

## Results

### Drought effect on leaf and stem physiology

Drought had a severe effect on water relations of lemon trees. The number of embolised conduits observed in microCT scans increased following drought (Fig. 2). Drought decreased hydraulic conductivity of control lemon trees (40 ppm K^+^, 400 ppm CO_2_) about ten-fold compared to pre-drought conditions (Fig. 2; Table 1). Leaf water potential decreased gradually in pine, and more so in lemon (Fig. 3). Drought effect on embolism level was also evident across potassium treatments (Fig. 4). Carbon assimilation (A) and transpiration (E) were reduced by about 50% and 70% (respectively) by drought (Fig. 2; Fig. S1). In pines, drought affected leaf gas exchange dramatically, decreasing E in the control trees by 90% (Fig. 2) and turning A negative, i.e. respiration rate surpassed assimilation rate (Fig. S1). However, the hydraulic conductivity of control pines decreased by 30% only (Fig. 2).

**Fig. 2.**
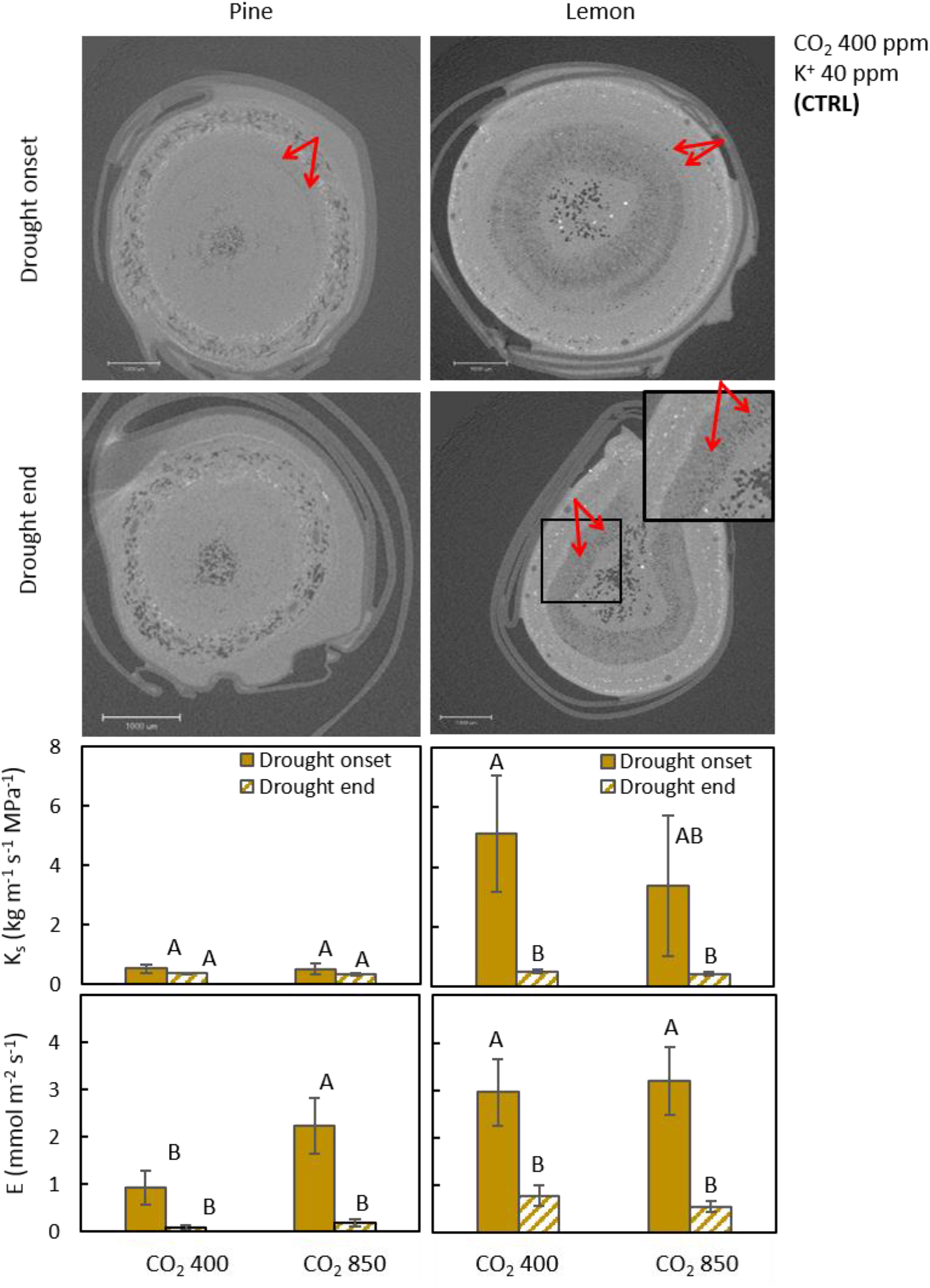
Images show cross-sections of pine (left) and lemon (right) branches wrapped in parafilm, taken from microCT scans (n=1). Black areas represent air, dark areas represent water or wood tissues, and white areas represent dense materials such as mineral or meristematic tissues. Inserts are enlarged portions of the sections, arrows point to embolised vessels. Size bars show 1000 μm. The lower panels show hydraulic conductivity (K_s_) and transpiration (E) of pine (left) and lemon (right) trees. Error bars represent standard error (n=3).

**Fig. 3.**
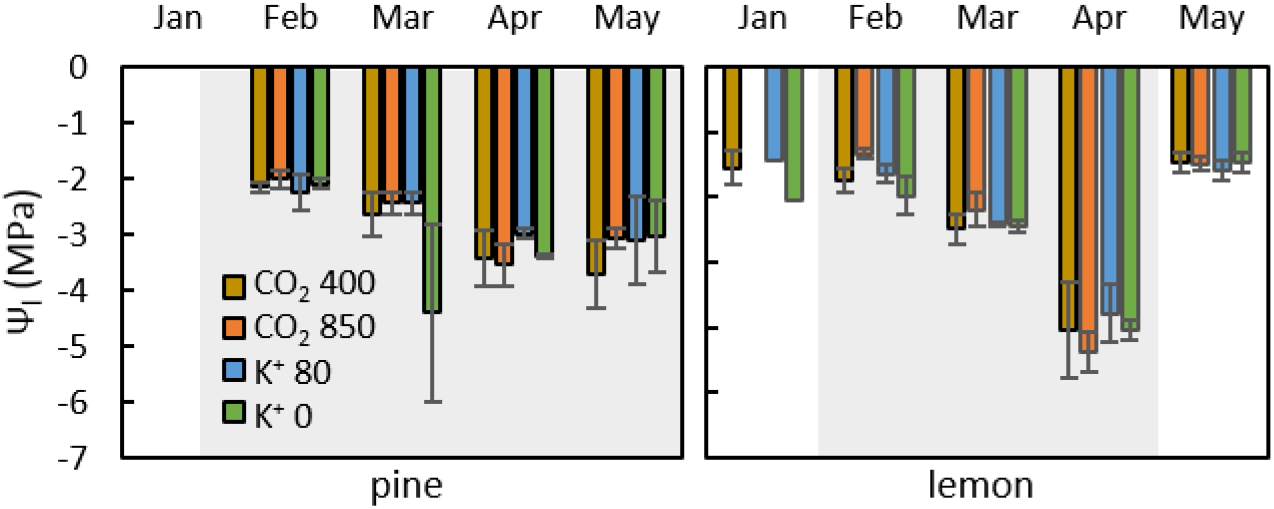
Leaf water potential (Ψ_1_) of pine (left) and lemon (right) trees treated with 0, 40, and 80 ppm K^+^ at 400 ppm CO_2_, and with 40 ppm K^+^ at 850 ppm CO_2_. Grey background mark drought period. Pre-drought measurements were done on lemon trees only. Error bars represent standard error (*n* = 1-5). No statistically significant difference was found between treatments within the same date nor between dates within the same treatment, according to Student’s *t* test (α=0.05).

**Fig. 4.**
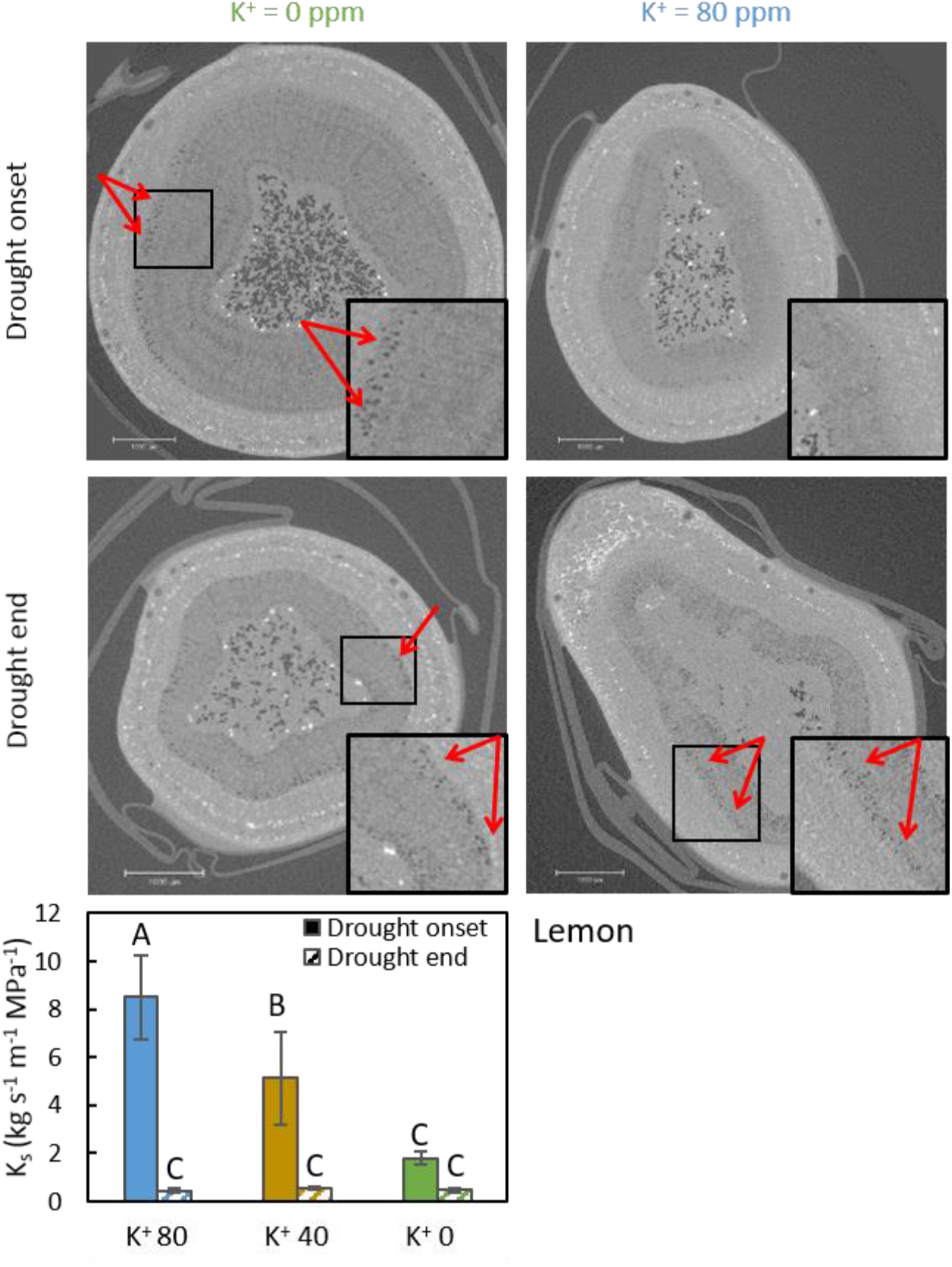
Images show cross-sections of branches from lemon trees treated with 0 ppm (left) and 80 ppm (right) K^+^ wrapped in parafilm, taken from microCT scans (n=1). Black areas represent air, dark areas represent water or wood tissues, and white areas represent dense materials such as mineral or meristematic tissues. Inserts are enlarged portions of the sections, arrows point to embolised vessels. Size bars show 1000 μm. The lower panel shows hydraulic conductivity (K_s_) of lemon trees treated with 0, 40 and 80 ppm K^+^. Error bars represent standard error (n=3)

### CO_2_ effect on leaf and stem physiology

Lemon trees at elevated CO_2_ had slightly less negative Ψ_1_ at the end of the pre-drought period than control trees (Fig. 3), but the effect was not statistically significant and was eliminated as drought progressed. Elevated CO_2_ did not induce any significant change in E and K_s_ in lemon trees, neither before nor after the drought period (Fig. 2). Interestingly, pine trees treated with elevated atmospheric CO_2_ had higher E values (2.23 mmol m^−2^ s^−1^) compared to the ambient CO_2_ pines (0.93 mmol m^−2^ s^−1^) at the end of the CO_2_ treatment (Fig. 2; significant by t-test but only at α = 0.10 level by ANOVA, Table1). This effect was not observed at the end of the drought (0.17 and 0.08 mmol m^−2^ s^−1^ for elevated and ambient CO_2_ pine trees, respectively). Additionally, there was no CO_2_ effect on pine hydraulic conductivity, Ψ_1_ or embolism levels (Figs. 2, 3).

### Potassium effect on leaf and stem physiology

Hydraulic conductivity increased gradually with increasing K^+^ concentration in the inert media in both lemon (Fig. 4; Table 1) and pine (Fig. 5; Table 1) before the drought. In lemon, almost no embolised vessels were observed in trees that were treated with 80 ppm K^+^. By the end of drought, though, K_s_ was low and similar among K^+^ treatments, and vessels were embolised even at high K^+^ (Fig. 4). No K^+^ effect was found on K_s_ nor on embolism rate in pine (Fig. 5).

**Fig. 5.**
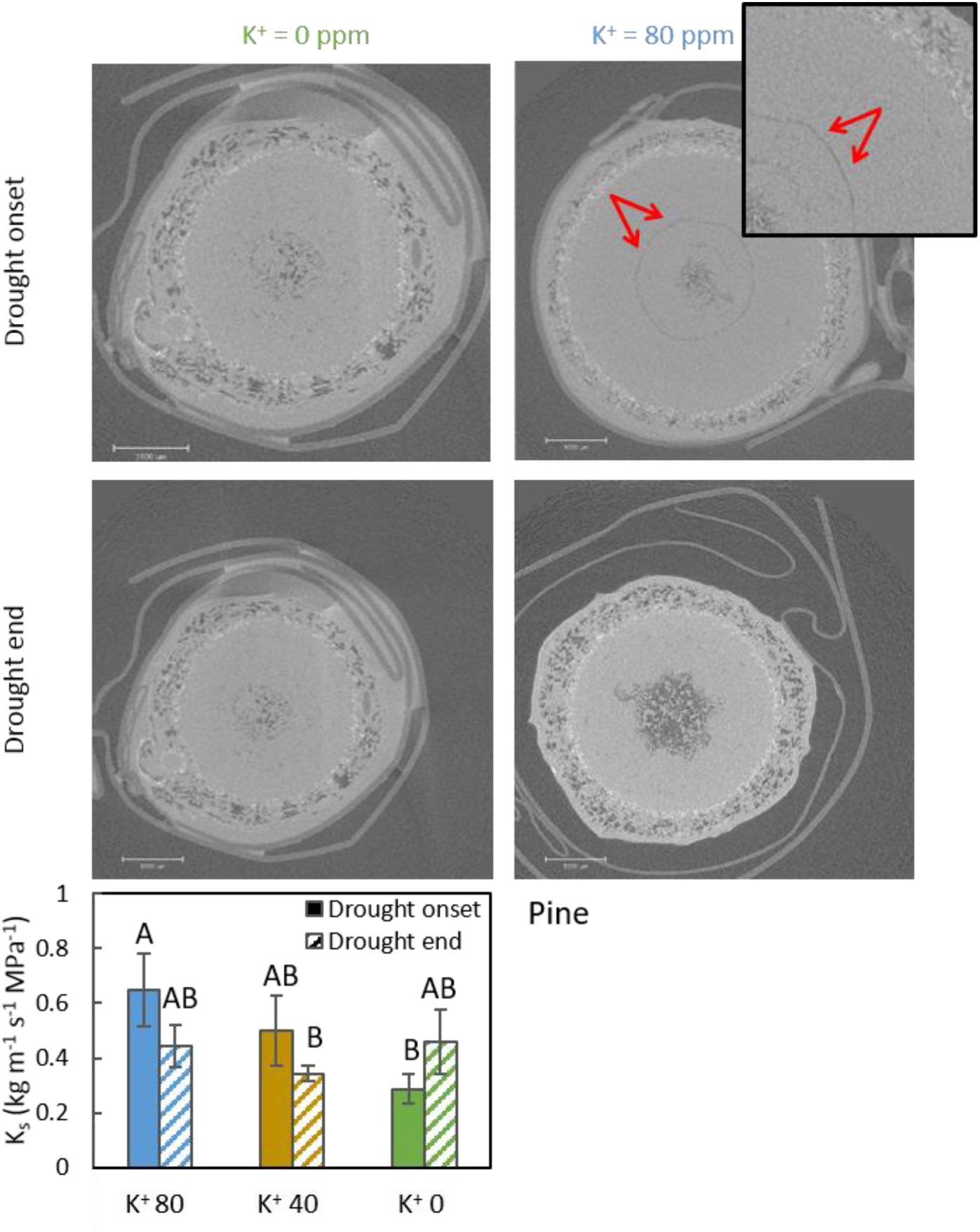
Images show cross-sections of branches from pine trees treated with 0 ppm (left) and 80 ppm (right) K^+^ wrapped in parafilm, taken from microCT scans (n=1). Black areas represent air, dark areas represent water or wood tissues, and white areas represent dense materials such as mineral or meristematic tissues. Inserts are enlarged portions of the sections, arrows point to embolised conduits. Size bars show 1000 μm. The lower panel shows hydraulic conductivity (K_s_) of pine trees treated with 0, 40 and 80 ppm K^+^. Error bars represent standard error (n=3).

## Discussion

Our study examined the effect of elevated K^+^ fertigation and atmospheric CO_2_ on the hydraulic response of young pine (conifer) and lemon (angiosperm) trees. The results suggest a strong effect of K^+^ on hydraulic conductivity in lemon and a weaker effect on pine, both of which comply with current explanations for the so-called ‘ion effect’, presumably due to different structure and function of the pits in the two functional groups. In this respect, our first hypothesis is partially confirmed. However, the response did not change the state of the trees by the end of drought. Elevated atmospheric CO_2_ seemed to affect transpiration in pines (but not in lemons), with the effect vanishing as well at the end of drought period, possibly due to the short-term exposure of the trees to elevated CO_2_. Our second hypothesis, on sugars assisting recovery from embolism, was hence disproved in our study system.

### Effects of potassium in the media on xylem conductivity and embolism

Previous studies reported a positive effect of xylem sap K^+^ levels on hydraulic conductivity, mainly in angiosperms (Van Ieperen et al. 2000; Zwieniecki et al. 2001, 2003; Lopes-Portillo et al. 2005; Trifilo et al. 2008; Aasamaa and Sõber 2010; Jansen et al. 2011; Nardini et al. 2011). However, Oddo et al. (2014) found no effect of K^+^ levels on the recovery from drought in laurel (*Laurus nobilis*) trees. In our study, lemon trees that were treated with increased K^+^ presented higher K_s_ compared to trees that received moderate or zero K^+^. To a lesser degree, the same phenomenon was observed in pines. This elevated K_s_ was in spite of the fact that all hydraulic measurements used the same, 19 mM KCl solution for all the trees. This implies that the excess K^+^ was not accumulated in the sap fluid, which would have been washed out during the measurements, but rather acted in living cells or was attached to the xylem conduits themselves

The difference between species in their hydraulic response to K^+^ may result from the structural difference in xylem pits. K^+^ is thought to affect hydraulic conductivity either through changes in the pectin matrix in the pit membrane (Lee et al. 2012; Zwieniecki and Secchi 2012) or through contrasting water potential and electrical gradients from opposite sides of the pit membrane (Santiago et al. 2013). In both cases, the different structure and function of pits in conifers would presumably be less affected by K^+^ levels, since torus margo acts more as a seal compared to the permeable nature of angiosperms pit membrane (Hacke et al. 2004; Cochard et al. 2009; Delzon et al. 2010). Interestingly, the effect on lemons was shown not only in hydraulic conductivity, but also in micro CT scans, which represent embolism levels. According to a common hypothesis, the increase in hydraulic conductivity under high K^+^ levels stems from enhanced hydraulic conductivity of non-embolised vessels, and therefore should not affect embolism *per se* in the xylem (Trifilò et al. 2011). However, it is possible that higher hydraulic conductivity increased the available water in the xylem tissue throughout the tree, perhaps via increased hydraulic permeability of the roots, thereby preventing cavitation events. In consistence with Oddo et al. (2014), the K^+^ effect on K_s_ under high water availability had no continuing impact on the hydraulic state of the trees after drought. This complies with both hypothesized K^+^ effects on pit membrane described above. Under low water availability, increasing the conductivity of pit membranes in active vessels would not increase the overall hydraulic conductivity, because water availability, rather than pit resistivity, is the limiting factor to conductivity under these conditions. If anything, high permeability of the pit membrane would increase its susceptibility to air seeding and to the spread of embolism through the xylem tissue (Tyree and Sperry 1989), thus reducing K_s_. An alternative explanation would be that fluctuations in K^+^ concentration in the sap are rapid, and the excess ions are quickly removed from the system. In such a scenario, both pre- and post-drought exposure to high potassium is futile in respect of the xylem water transport.

### CO_2_ effect on leaf and stem physiology

We did not observe any effect of elevated CO_2_ on the physiology of lemon trees, namely on A. Therefore, it is likely that the CO_2_ treatment did not increase the trees’ carbohydrate reservoirs. Previous studies on the impact of elevated CO_2_ on plant physiology mostly described an increased carbon assimilation and reduced stomatal conductance (Medlyn et al. 1999; Ainsworth and Long 2004; Leakey et al. 2009; Pazzagli et al. 2016; Lotfiomran et al. 2016). However, Curtis and Wang (1998) found increased assimilation under elevated CO_2_ with no effect on stomatal conductance, and Phillips et al. (2011) found stomatal conductance to be affected by CO_2_ level only under elevated temperature. Additionally, other studies have shown an effect of elevated CO_2_ on growth (Paudel et al. 2018) or on nocturnal water loss (Zeppel et al. 2011, 2012, 2014)_only when combined with drought. Medlyn et al. (2001) found in a meta-analysis study contradicting results regarding assimilation response to elevated CO_2_ and concluded that the length of CO_2_ treatment is highly important to induce a response, especially in woody species. This conclusion is also in agreement with the lack of response found in our study, as the elevated CO_2_ treatment was only gives for 30 days period. The increased transpiration of pines under elevated CO_2_ (significant by t-test, Fig. 2, and *P* = 0.088 by ANOVA, Table 1) contrasts with the commonly observed transpiration response to high CO_2_ (Morgan et al. 2004). Higher water loss would call for quicker dehydration of the media and a more severe effect of drought on these trees, yet no such trend was observed. These data may imply high stomatal sensitivity to environmental changes. If this is the case, then the pines that lost more water due to elevated transpiration would later, when exposed to drought, compensate for that loss with earlier stomatal closure. In order to test for the validity of that hypothesis, measurements would have to be conducted in a way that captures the dynamics of physiological behavior as drought progresses.

### Study limitations

During the experiment, several methodological issues were encountered. A major issue was the inability to conduct hydraulic measurements on intact samples. In addition, due to logistical constraints, the CO_2_ treatment was only imposed for 30 days. Embolism sampling has long been a concern in plant hydraulic research. When cutting a branch that transports water under tension, air bubbles are introduced into the system and create artificial embolism (Dixon 1914; Wheeler et al. 2013; Sperry 2013; Rockwell et al. 2014; Torres-Ruiz et al. 2015). Additionally, when vessel length exceeds the length of measured branch segment, the resistance posed on water flow by the xylem tissue is eliminated and the vessel is rapidly drained, deeming the hydraulic conductivity measurements not valid (Cochard et al. 2005, 2010). These artefacts are less relevant for measurements on conifers, due to the short length of tracheid elements (Sperry et al. 2006). However, they pose a bigger problem when long-vesseled species are measured. Several protocols were designed with the purpose of overcoming this challenge (Wheeler et al. 2013; Torres-Ruiz et al. 2015; Klein et al. 2018; López et al. 2018) mostly adjusting the sample size, cutting and preparation procedures. As stated in the methods sections, in our study, all precautions were taken during sampling, and comparisons were only conducted within species in order to avoid biased results. Additionally, data regarding vessel length of *C. limon* were examined, and the percentage of opened vessels in the measured segments was found to be negligible (see methods). Additionally, since the methods used here to evaluate hydraulic conductivity were destructive, in each measurement a single branch was used as a representative for all branches on the tree, assuming similarity in hydraulic state between all branches. When technical conditions allow it, repeating measurements on the same sample may shed a light on the validity of this methodology. Other limitations include the short-term duration that trees were exposed to CO_2_, only 30 days of their five year life span. The duration that trees are exposed to CO_2_ can impact on whether water relations and tree physiology are altered or not (Medlyn et al. 2001, 2011). Finally, due to the short-termed and subtle effects described in this study, it seems that in order to capture the complete effect of treatments on the parameters measured here, the measurements should be conducted under a tighter temporal scale. For example, understanding the complete dynamics of K^+^ effect may require weekly measurements of hydraulic conductivity, leaf water potential and sap composition.

### Implication for orchard and forest tree physiology

Our results show a positive linear effect of potassium on tree hydraulic conductivity, with a stronger effect on lemon compared to pine. This result strengthens the hypothesis of action through changes in the pectin matrix in the angiosperm pit membrane, with lesser implications for gymnosperms. Furthermore, the unexpected effect of high potassium reducing xylem native embolism (at mesic conditions) suggests a potential relevance for tree health. However, the short-termed effect of potassium on xylem hydraulic conductivity questions the relevance of these parameters to trees’ survival under drought. Instead, it might be that high potassium level can be important under normal growth conditions. The positive, additive effect of potassium concentration occurred at both leaf and xylem levels, and might contribute to the long-term health status of broadleaf trees in general, and particularly to evergreen broadleaf trees grown in orchards. More research is needed to test whether these benefits can be translated into higher growth or yield, and if potential tradeoffs exist. Considering the ever growing demand for fresh fruits globally, even a small enhancement can be important.

## Acknowledgments

We thank Eran Raveh, Uri Yermiyahu and Moshe Halpern (ARO Volcani in Gilat research station) for their significant contribution to pre-drought treatments designing and execution. Special thanks go to the team of Prof. Ari Schaffer (ARO Volcani in Beit Dagan) for HPLC analyses of sap sugars. The manuscript benefitted significantly from useful comments and suggestions made by Dr. Melanie Zeppel (Macquarie University, NSW, Australia) and Prof. Andrea Nardini (University of Trieste, Italy). TK wishes to thank the Benoziyo Fund for the Advancement of Science; Mr. and Mrs. Norman Reiser, together with the Weizmann Center for New Scientists; and the Edith & Nathan Goldberg Career Development Chair.

## Authors’ contributions

YW conducted all experiment and measurements, and wrote the manuscript.

VB assisted with microCT scans and analysis, and edited the manuscript.

JMG assisted with statistical analysis and edited the manuscript.

TK initiated the research, supervised all stages of the study and edited the manuscript.

## Supplementary Information

**Table. S1.** Fertilizer composition for the three K^+^ treatments: concentrations of compounds used in each formula (in mmol, top) and the resulting ion concentrations (in ppm, bottom). pH was adjusted at the range of 5-5.7 across the three treatments. All fertilizer formulas had the same number of anions and cations except for K^+^ and Cl^−^

| (mmol) | 0 ppm $k^+$ | 40 ppm $K^+$ | 80 ppm $K^+$ |
| --- | --- | --- | --- |
| $NH_4NO_3$ | 0.055 | 0.055 | 0.055 |
| $Ca(NO_3)_2$ | 2.51 | 2.51 | 2.51 |
| $MgCl_2$ | 1.0 | 1.0 | 1.0 |
| KCl | 0 | 1.0 | 2.04 |
| $NH_4H_2PO_4$ | 0.3 | 0.3 | 0.3 |

| ppm | 0 ppm $k^+$ | 40 ppm $K^+$ | 80 ppm $K^+$ |
| --- | --- | --- | --- |
| $NH_4^+$ | 5 | 5 | 5 |
| $NO_3^-$ | 36 | 36 | 36 |
| P | 9.5 | 9.5 | 9.5 |
| $K^+$ | 0 | 40 | 80 |
| $Mg^{2+}$ | 24.3 | 24.3 | 24.3 |
| $Ca^{2+}$ | 120 | 120 | 120 |

**Fig. S1.**
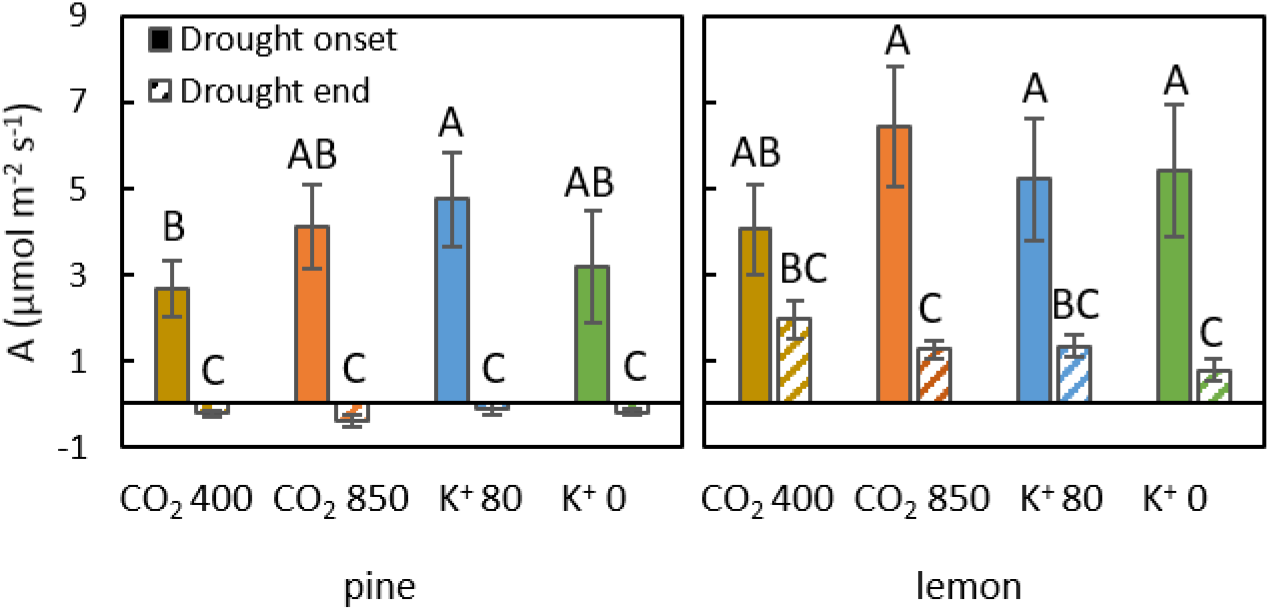
Carbon assimilation rates of pine (left) and lemon (right) trees measured at drought onset (full columns) and drought end (striped columns). Error bars represent standard error (n=3) and different letters represent significant difference between values according to Student’s *t* test (α=0.05)

## Notes

### Competing Interest Statement

The authors have declared no competing interest.

